# Naproxen impairs load-induced bone formation, reduces bone toughness, and delays stress fracture repair in mice

**DOI:** 10.1101/427138

**Authors:** Jino Park, Andrzej Fertala, Ryan E. Tomlinson

## Abstract

Debilitating stress fractures are surprisingly common in physically active individuals, including athletes, military recruits, and dancers. These individuals are overrepresented in the 30 million daily users of non-steroidal anti-inflammatory drugs (NSAIDs). We hypothesized that regular use of NSAIDs would predispose habitually loaded bones to stress fracture and delay the repair of these injuries. To test this hypothesis, adult mice were subjected to six bouts of axial forelimb compression over two weeks. Aspirin, naproxen, or vehicle was administered 24 hours before loading. Naproxen-treated mice had diminished load-induced bone formation as well as a significant loss in toughness in non-loaded bone, which were not observed in aspirin-treated mice. Furthermore, there were no differences in RANKL/OPG ratio or cortical bone parameters. Picrosirius red staining and second harmonic generation imaging revealed that alterations in bone collagen fibril size and organization were driving the loss of toughness in naproxen-treated mice. Separately, adult mice were subjected to an ulnar stress fracture generated by a single bout of fatigue loading, with NSAIDs provided 24 hours before injury. Both aspirin-treated and naproxen-treated mice had normal forelimb use in the week after injury, whereas control mice favored the injured forelimb until day 7. However, woven bone volume was only significantly impaired by naproxen. Both NSAIDs were found to significantly inhibit *Cox2* and *Ngf* expression following stress fracture, but only naproxen significantly affected serum PGE2 concentration. Overall, our results suggest that naproxen, but not aspirin, may increase the risk of stress fracture and extend the healing time of these injuries, warranting further clinical evaluation for patients at risk for fatigue injuries.

## INTRODUCTION

One of the principal roles of the mammalian skeleton is to sustain the mechanical loads necessary for terrestrial function and movement; fracture occurs when these applied loads exceed the bone strength. However, bone tissue is also susceptible to fatigue, a process by which repetitive loads that do not cause fracture lead to the formation and propagation of cracks in bone. These painful and debilitating injuries, known as stress fractures, generally require the complete cessation of activity for weeks or months to allow the bone to repair itself [1] and may necessitate surgical stabilization [2]. Stress fractures are particularly common in soldiers, athletes, and performers, with documented incidence rates up to 50% per year [3-6]. Nonetheless, since treatment of these injuries represents a significant economic burden with few clinical options, identifying stress fracture risk factors other than physical activity is an important orthopaedic research goal.

Non-steroidal anti-inflammatory drugs (NSAIDs) are the most commonly consumed medication in the world, with over 30 million daily users [7,8]. Regular NSAID users include as many as 50% of patients over the age of 65 [9] and 80% of active duty US military [10]. NSAIDs are effective pain relievers by blocking the cyclooxygenase (COX) enzyme isoforms, COX1 and COX2, which prevents the synthesis of prostaglandin H2 (PGH2), the precursor to downstream prostaglandins (PGE2, PGF2α, PGD2) as well as thromboxane (TXA2) and prostacyclin (PGI2) [8,11]. Whereas the constitutively active COX1 generates prostaglandins that are required to maintain physiological functions, such as gastric cytoprotection and platelet aggregation, COX2 produces prostaglandins that drive fever and inflammation [12]. COX2-specific inhibitors, such as celecoxib, were introduced to minimize the gastrointestinal bleeding and platelet abnormalities associated with non-specific NSAIDs [13,14]. Results from the 2016 PRECISION trial relieved concerns regarding the cardiovascular complications of celecoxib, opening a potential floodgate for increased prescription of COX2-specific NSAIDs [15].

Inhibition of COX2 enzymatic activity by any NSAID decreases PGE2 synthesis. However, PGE2 is part of an inflammatory signaling pathway that is critical for load-induced bone formation [16-18]. In general, bone is formed in response to mechanical forces at the sites of highest strain and removed in the areas of lowest strain [19]. This process, referred to as strain adaptive bone remodeling, enables bone to efficiently adapt to functional demands by generating bone where it is needed and eliminating bone that is underutilized – a process that has been shown to greatly increase the fatigue strength of bone [20]. Recent work by Hughes et al. reported that any NSAID prescription was associated with a significantly increased risk for stress fracture in US Army Soldiers, and the risk increased substantially (RR: 5.3, 95% CI: 4.9-5.7) for Soldiers during Basic Combat Training, a period of intense physical activity [21]. In that study, naproxen was the second most commonly prescribed NSAID and was found to be associated with the greatest increase in risk. They attributed the increase in stress fracture risk to diminished load-induced bone formation, which has been previously demonstrated following NSAID administration in clinical and preclinical studies [22-26], but this explanation may not be sufficient to explain the surprisingly large increase in risk or stratification by NSAID type.

To dissect the mechanism by which different NSAIDs may affect the skeletal response to mechanical forces, we focused on two popular NSAIDs: naproxen and aspirin. We hypothesized that aspirin, a relatively poor COX2 inhibitor with newly identified targets outside of prostaglandin signaling [27,28], may be less deleterious to bone than naproxen, a potent COX2 inhibitor [29]. To test this hypothesis, we determined the effects of clinically relevant dosages of aspirin and naproxen on load-induced bone formation and stress fracture repair in mice. Our results reveal multiple mechanisms by which naproxen, but not aspirin, may increase the risk and diminish the repair of fatigue injuries.

## RESULTS

### Naproxen, but not aspirin, impaired load-induced bone formation

Normal strain adaptive bone remodeling is critical for increasing fatigue resistance of bones. To test if reasonable doses of NSAIDs would negatively affect load-induced bone formation, adult female C57BL/6J mice were subjected to six bouts of axial forelimb compression (3 N, 100 cycles) over a period of two weeks, starting at 16 weeks of age. Drinking water was used to administer aspirin (100 mg/kg/d), naproxen sodium (10.9 mg/kg/d), or vehicle (water) starting 24 hours before loading. Calcein and alizarin were administered 3 and 10 days after the first bout of loading, respectively, and could be readily visualized in PMMA-embedded sections (Figure 1A), which were analyzed quantitatively by dynamic histomorphometry (Table S1). In vehicle treated mice, axial compression produced the expected anabolic response in loaded limbs, with robust increases in bone formation evident on the periosteal and endosteal surfaces of loaded ulnae. Consistent with its characterization as a relatively weak COX2 inhibitor, administration of aspirin did not significantly affect bone formation parameters. However, mice treated with naproxen had significantly decreased relative (loaded – non-loaded) periosteal bone formation parameters (Figure 1B-D), including decreases in mineralizing surface per bone surface (−65%), mineral apposition rate (−77%), and bone formation rate per bone surface (−76%). There were no significant differences in endosteal bone formation parameters (Figure 1E-G).

**Figure 1.**
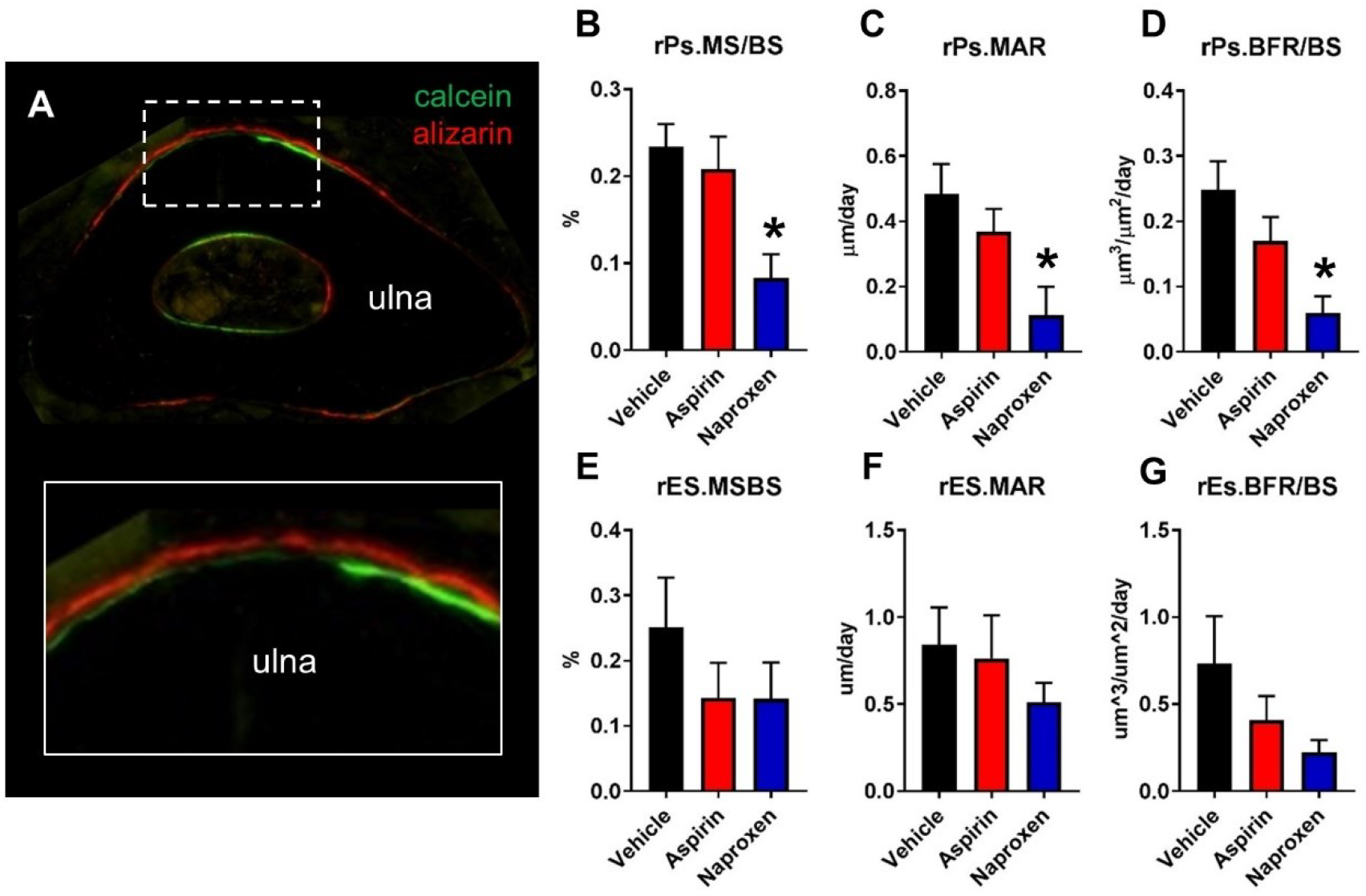
Naproxen treatment diminished load-induced bone formation by dynamic histomorphometry. Bone formation was quantified by dynamic histomorphometry using A) ulnar cross sections with alizarin (red) and calcein (green) bone formation labels, with outcomes including B) relative periosteal mineralizing surface per bone surface (rPs.MS/BS), C) relative periosteal mineral apposition rate (rPs.MAR), and D) relative periosteal bone formation rate per bone surface (rPS.BFR/BS). * p < 0.05 vs. vehicle, scale bars are 100 microns. n = 6-7 per group.

### Treatment with naproxen, but not aspirin, resulted in significant loss of femoral toughness

Determining the systemic effects of NSAIDs on overall bone remodeling is critical for interpreting our findings regarding load-induced bone formation. Therefore, we analyzed the serum markers RANKL and OPG. Consistent with our overall hypothesis, RANKL/OPG ratios were not affected by NSAID treatment (Figure 2A). In addition, we performed microCT and three-point bending analysis of the femur – a bone not subjected to additional mechanical load by forelimb axial compression. As expected, we found no significant differences in bone volume, cortical thickness, or bone mineral density between treatment groups by microCT (Figure 2B-D). In addition, ultimate moment, ultimate stress, bending rigidity, and Young’s modulus – structural and material properties that primarily define elastic behavior – were not affected by NSAID treatment (Figure 2E-H). Surprisingly, we observed a large and significant decrease (−36%) in the amount of energy required to fracture bones from mice treated with naproxen (Figure 2I), which was mainly due to decreased (−45%) post-yield energy (Figure 2J). After accounting for geometry, we still observed a significant decrease (−35%) in toughness in bones from mice treated with naproxen as compared to vehicle (Figure 2K), which was also mainly due to decreased (−45%) post-yield toughness (Figure 2L). The effect of aspirin on either toughness or energy was not statistically significant. Representative moment-displacement curves illustrate the striking alterations in post-yield behavior induced by naproxen, but not aspirin or vehicle (Figure S1). To further examine this phenomenon, we performed notched three-point bending. In this case, we found no difference in the critical stress intensity factor (KIC), a measure of resistance to crack initiation at a dominant flaw (Figure 2M).

**Figure 2.**
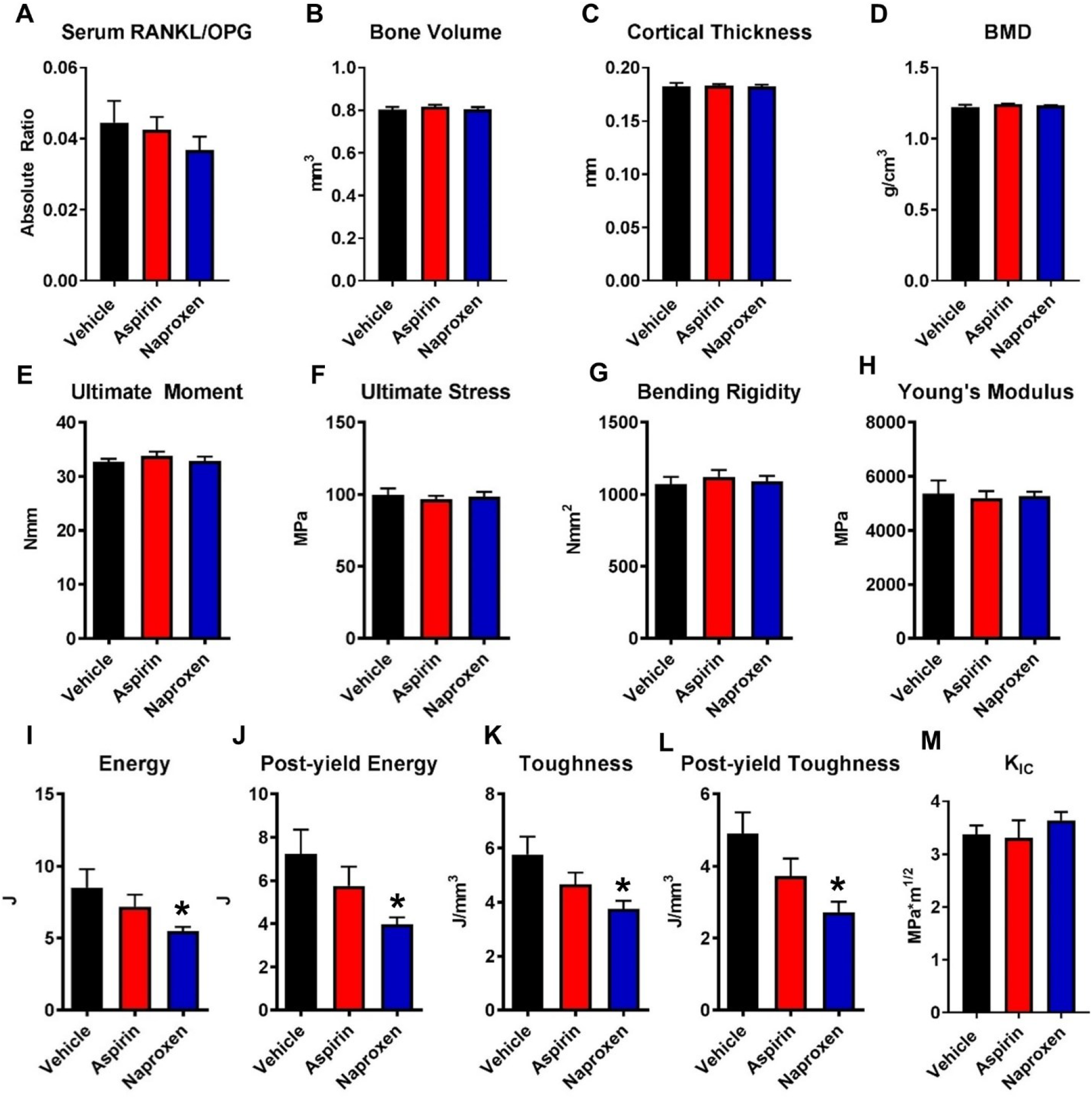
Effects of aspirin and naproxen on non-loaded bones. A) RANKL/OPG ratio was quantified by ELISA. MicroCT was used to quantify B) Bone volume, C) Cortical Thickness, and D) Bone Mineral Density in the femur. Standard three-point bending of the femur was used to determine E) Ultimate Moment, F) Ultimate Stress, G) Bending Rigidity, H) Young’s Modulus, I) Energy to Failure, J) Post-yield energy, K) Toughness, and L) Post-yield toughness. M) Notched three-point bending was used to determine KIC.* p < 0.05 vs. vehicle, n = 6-7 per group.

### Aspirin and naproxen affect collagen content and structure in bone

Since bone derives much of its toughness from its organic phase [30], we attempted to determine a potential mechanism for the loss of toughness observed in bones from naproxen-treated mice by analyzing collagen content and structure in the tibia and femur. First, paraffin-embedded sections from the femoral midshaft stained with picrosirius red were visualized using bright-field (Figure 3A-C) and polarized light (Figure 3D-F). Next, these sections were visualized using second-harmonic generation (SHG) imaging to analyze fibrillar collagen (Figure 3G-L). Quantification of both imaging modalities revealed similar alterations in collagen structure and organization. Color thresholds, used to determined collagen fibril thickness in picrosirius red stained slides, revealed a significant increase (+201%) in the percentage of thin (green) fibrils in bones from naproxen treated mice as compared to vehicle (Figure 3M). This increase in thinner fibrils was at the expense of thick (red) fibrils, which were significantly decreased (−49%) in naproxen treated mice as compared to vehicle. There were no significant differences between aspirin treated mice and vehicle control. Consistent with these results, we found a significant decrease in SHG signal in the cortical bone in mice treated with naproxen (−32%), but there was no significant effect of aspirin (Figure 3N). To determine if these alterations were due to a net loss in bone collagen, we analyzed a sample of the mid-tibial diaphysis using a colorimetric assay for 4-hydroxyproline (a required component of the collagen helical structure). Here, treatment with either aspirin or naproxen significantly decreased collagen content in the tibia (Figure 3O), but neither NSAID significantly affected serum levels of 4-hydroxyproline (Figure 3P). However, collagen fractions extracted from tibial samples displayed no differences in collagen crosslinking between groups, as determined by SDS-PAGE (Figure 4A) with quantification (Figure 4B-E).

**Figure 3.**
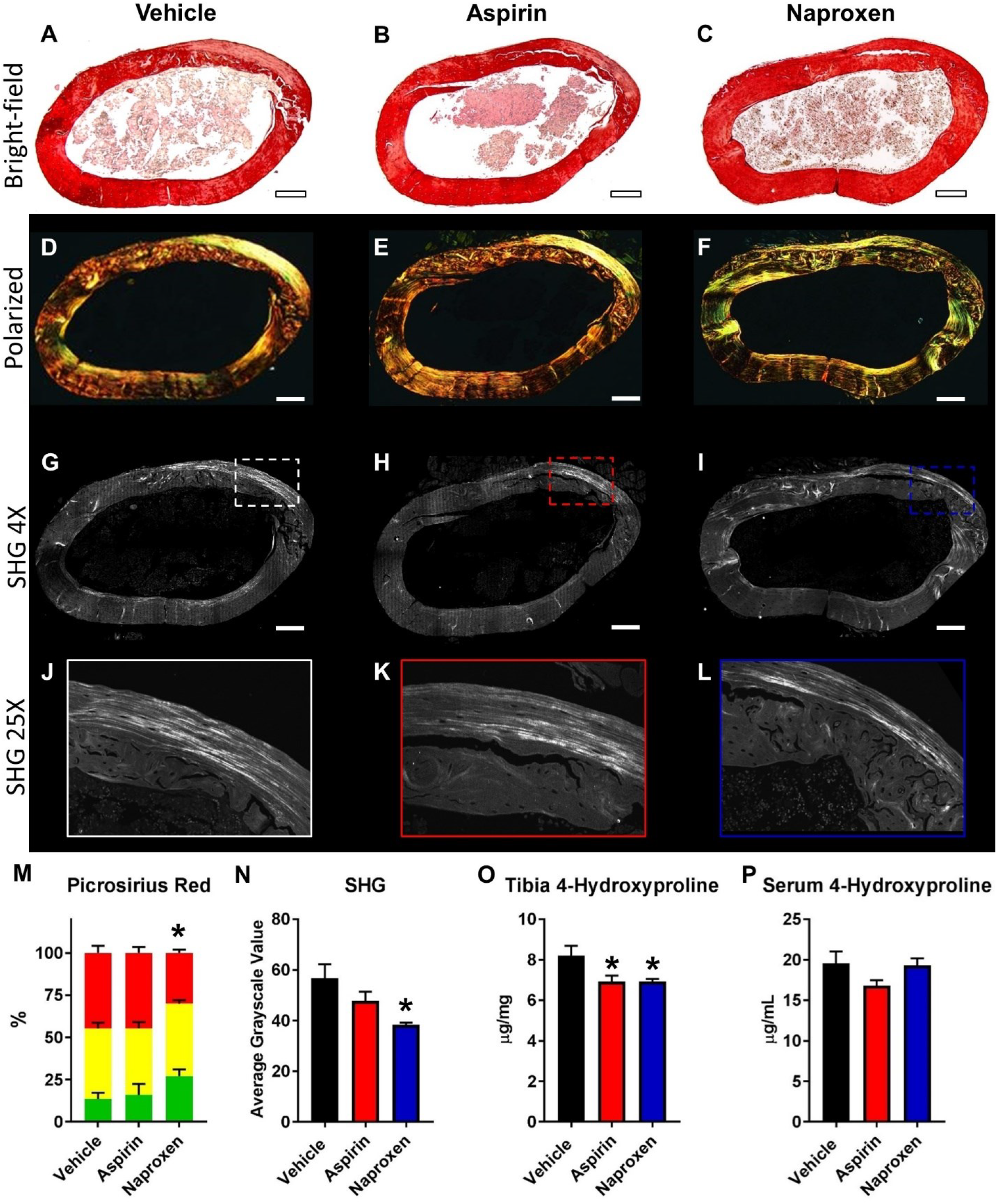
Effects of Aspirin and Naproxen on bone collagen content and organization. Quantification of 4-hydroxyproline in the A) tibia and B) serum of treated mice. C) Quantification of sections stained with picrosirius red and imaged using polarized light. D) Quantification of SHG signal in sections. E-J) Representative picrosirius red stained sections under E-G) bright-field and H-J) polarized light. K-P) Representative SHG images. * p < 0.05 vs. vehicle, scale bars are 100 microns. n = 6-7 per group.

**Figure 4.**
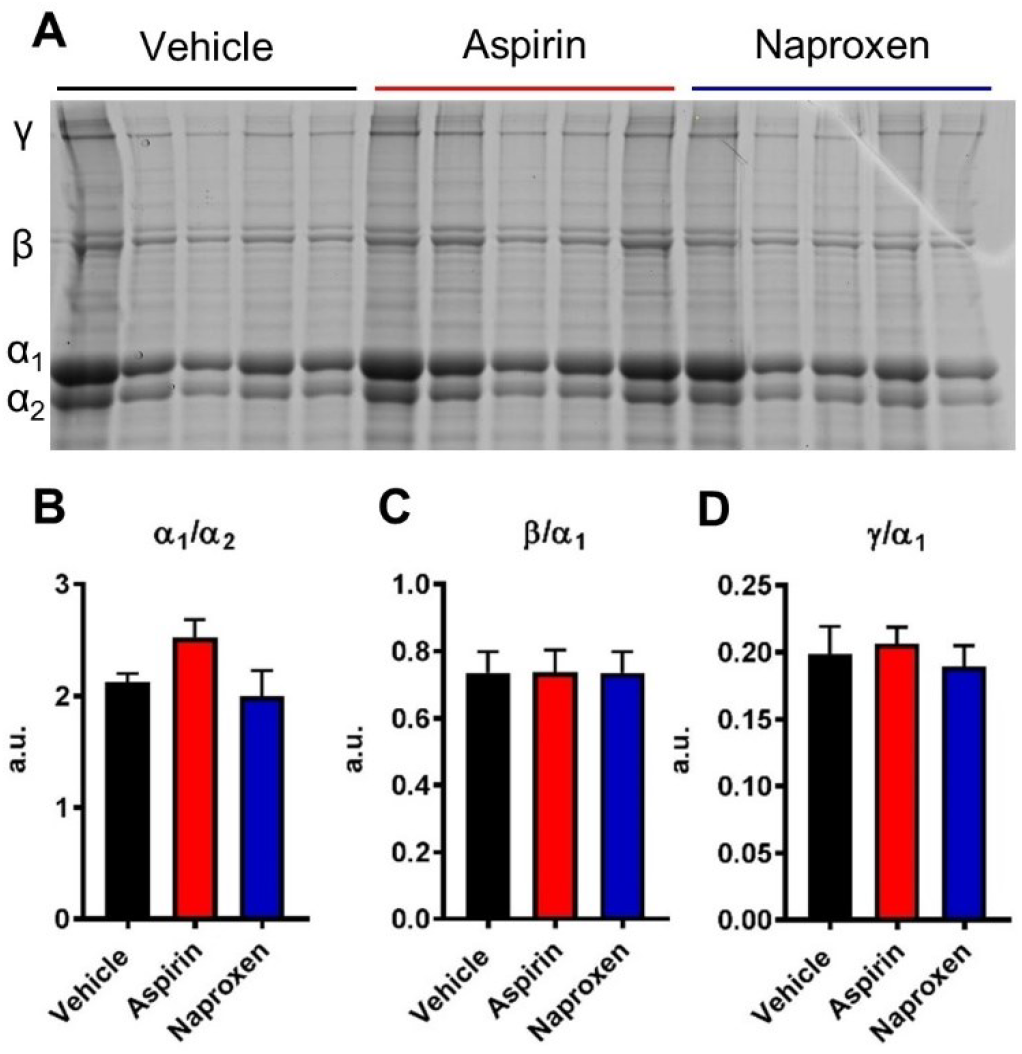
Crosslinking analysis of bone collagen. A) Collagen extracted from tibial samples was analyzed by SDS-PAGE with quantification of B) α_1_/ α_2_, C) β/α_1_, and D) γ/α_1_. n = 5 per group.

In total, these results suggest that stress fracture risk may be elevated by regular use of naproxen, due to a combination of decreased load-induced bone formation as well as diminished bone toughness resulting from alterations in collagen content, structure, and organization.

### Both aspirin and naproxen provide analgesia for stress fracture-related pain

NSAIDs are routinely initiated to treat stress fracture-related pain, but the effects on stress fracture healing are not clearly defined. To determine the specific effects of aspirin and naproxen in this scenario, we utilized a standard rodent preclinical model of stress fracture. Here, C57BL/6J mice were subjected to an ulnar stress fracture generated by a single bout of fatigue loading at 16 weeks of age. As before, drinking water containing aspirin (100 mg/kg/d), naproxen sodium (10.9 mg/kg/d), or vehicle was provided 24 hours before loading. To assess the efficacy of each NSAID to relieve stress fracture-related pain at the prescribed dose, forelimb asymmetry testing was performed one day before stress fracture, and then daily after stress fracture, starting 3 days following the procedure. We observed that mice receiving vehicle used their loaded forelimb significantly less (−25% to −34%) on days 3 to 6 as compared to baseline (Figure 5). By day 7, vehicle treated mice no longer used their loaded limb less than baseline. In contrast, there was no significant difference in forelimb usage in mice treated with either aspirin or naproxen at any time point. In total, these data indicate that this preclinical model of stress fracture is associated with discomfort resulting in reduced forelimb usage in the first 6 days after loading that can be alleviated by modest doses of either naproxen or aspirin.

**Figure 5.**
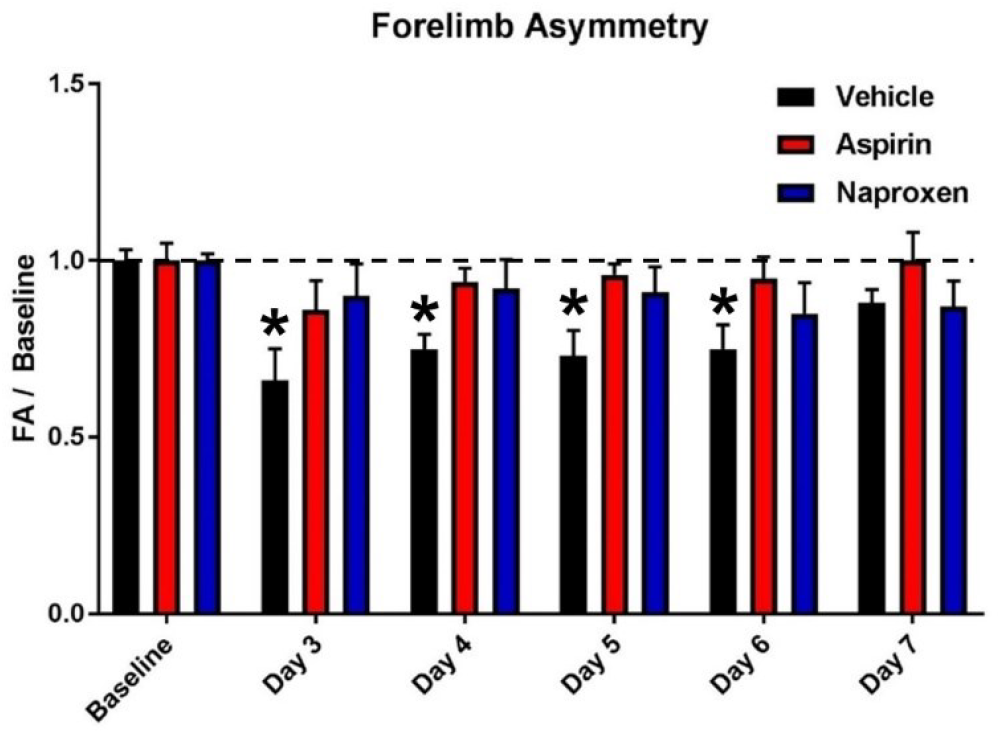
Effect of NSAIDs on forelimb usage after stress fracture. Forelimb usage scores were normalized to baseline for each group. * p < 0.05 vs. baseline by two-way ANOVA with repeated measures. n = 6-7 per group.

### Naproxen, but not aspirin, impairs stress fracture healing

To determine if treatment with these NSAIDs affected stress fracture repair, we harvested intact forelimbs from mice 7 days after stress fracture. In these loaded limbs, robust woven bone formation was observed at the ulnar midshaft in all mice (Figure 6A-C). Quantification of the crack length was not significantly different between groups, suggesting each animal received a similar amount of fatigue damage (Figure 6D). Similarly, the bone mineral density of the woven bone was not significantly different between groups (Figure 6E). However, the amount of woven bone formed in response to injury was significantly diminished (−27%) in bones harvested from mice treated with naproxen (Figure 6F). Consistent with this finding, we observed a trend (p = 0.09) that the woven bone in naproxen-treated mice tended to occupy less of the length of the ulna than in vehicle-treated mice, but this observation was not statistically significant between groups (Figure 6G). Furthermore, naproxen-treated mice had significantly decreased serum concentration of PGE2 (Figure 6H). However, we did not observation any alterations in serum RANKL/OPG ratio between treatment groups at this time point (Figure 6I). In total, these data indicate that naproxen treatment significantly diminishes stress fracture repair due to an effective blockade of COX2/PGE2 signaling, unlike treatment with aspirin.

**Figure 6.**
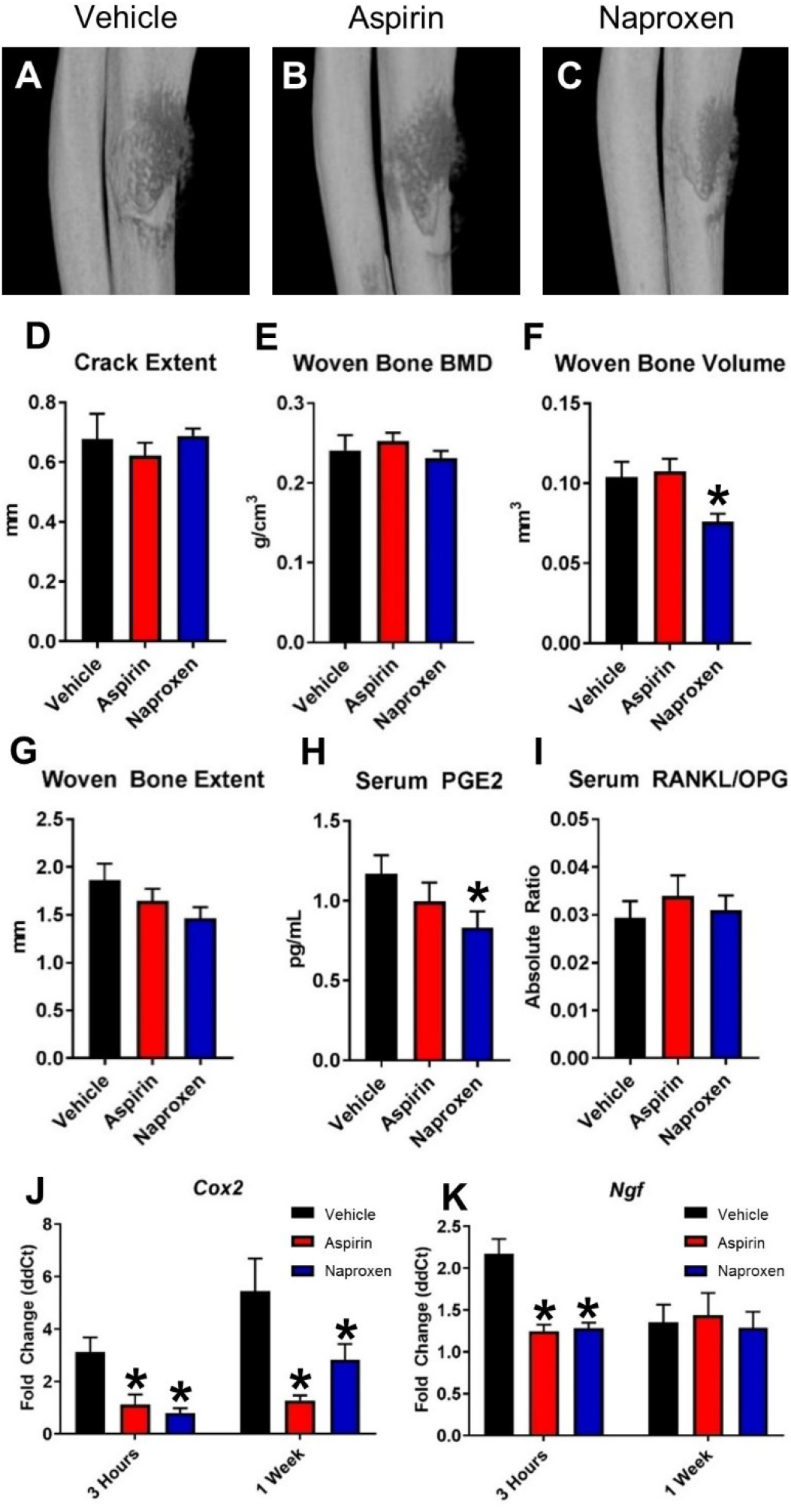
Naproxen significantly impairs stress fracture healing in mice. Representative microCT reconstructions of forelimbs from A) Vehicle B) Aspirin and C) Naproxen treated mice were quantified after 7 days of healing by D) Crack extent, E) Woven bone BMD, F) Woven Bone Volume, and G) Woven Bone Extent. Furthermore, serum was used to quantify H) PGE2 and I) RANKL/OPG ratio at this time point. mRNA harvested from the forelimb was used to quantify expression of J) Cox2 and K) Ngf. * p < 0.05 vs. vehicle, n = 5-7 per group.

### Aspirin and naproxen affect *Cox2* and *Ngf* expression following stress fracture

To further characterize the effect of NSAIDs on stress fracture healing, we isolated mRNA from the central third of the ulna in both loaded and non-loaded limbs. *Cox2* expression was assayed, as it plays a key role in the action of NSAIDs and is itself a target of PGE2 signaling. Three hours after stress fracture, *Cox2* expression was significantly increased in the loaded limbs of vehicle treated mice as compared to non-loaded limbs. In contrast, *Cox2* expression in loaded limbs from both aspirin and naproxen treated mice was not significantly different from non-loaded limbs and was significantly decreased as compared to vehicle treatment (Figure 6J). This pattern was similar after 7 days. In addition, we quantified the transcriptional activity of nerve growth factor (*Ngf*), which plays a central role in mediating osseous pain [31]. We found that *Ngf* expression was significantly upregulated at 3 hours in vehicle treated mice, similar to previous reports at 1 hour in rat [32]. Surprisingly, this increase in *Ngf* expression was entirely blocked by administration of either aspirin or naproxen (Figure 6K). We did not observe any increase in *Ngf* expression in any group 7 days after loading.

## DISCUSSION

In this study, we analyzed the effects of two popular NSAIDs, naproxen and aspirin, on load-induced bone formation and stress fracture repair in mice. Our main objectives were to determine if routine use of these drugs at recommended dosages would increase the risk of stress fracture during periods of intense physical activity or impair the healing process following a fatigue injury. In total, we observed that naproxen, but not aspirin, was associated with a significant decrease in load-induced bone formation and an unexpected loss of bone toughness over a period of two weeks of loading. Furthermore, naproxen, but not aspirin, was associated with significantly diminished woven bone formation following stress fracture, although both relieved stress fracture-related pain equally well. As a result, we conclude that the routine use of naproxen may increase the risk of stress fracture in active individuals and/or extend the time required for healing, and therefore warrants further clinical investigation and cautious use in subjects with routine intense physical activities.

In this study, we demonstrated that naproxen, but not aspirin, significantly decreased bone formation induced by axial forelimb compression in mice using standard dynamic histomorphometric measurements. This result is consistent with previous reports showing that NSAIDs (primarily indomethacin) have a strong negative effect on strain adaptive bone remodeling in rodents [22-25]. In particular, a high dose of indomethacin administered before a single bout of loading, but not afterwards, has been shown to nearly completely block load-induced bone formation [23]. Our continuous administration of NSAIDs in drinking water over a two week period of loading essentially replicated this scenario, albeit with drug administration well below the maximum recommended dose. We anticipated that the potential risk of chronic NSAID usage to future stress fracture would be directly related to the response to mechanical loading, which greatly increases fatigue resistance with only modest increases in bone mass [20]. However, we were surprised to find that administration of naproxen resulted in a significant decrease in bone toughness that may drive increases in stress fracture risk, without differences in bone size, strength, elastic modulus, or mineral content. The decrease in toughness was nearly exclusively due to diminished post-yield deformation, which we attributed to alterations in bone collagen size and organization. We did not observe any changes in KIC (fracture toughness) as quantified using notched three-point bending of mouse femur. Calculating JC, which incorporates the contribution from plastic deformation, may have yielded additional insight given the alterations in post-yield behavior, but the analytical solution is not applicable to this testing scenario [33]. In addition, we did not consider other factors that could alter bone toughness, such as water and GAG content [34-36]. Furthermore, naproxen may have direct drug-bone effects, such as those recently described for raloxifene [37]. In total, more study is required to determine if the loss of bone toughness is a unique feature of naproxen or common to NSAIDs as well as the specific mechanism of action.

In contrast to naproxen, aspirin had no statistically significant effects on load-induced bone formation, bone toughness, or stress fracture repair in this study. A potential explanation for our results is that aspirin is rapidly converted to salicylic acid (SA) in the body, with a half-life of less than 20 minutes [38]. Although SA is a poor inhibitor of prostaglandin synthesis (more than 15-fold less potent than aspirin on a weight basis [27]), SA is widely used to treat pain, fever, and inflammation with essentially the same pharmacological effects as aspirin [39,40]. We attempted to compensate for the short circulation time of aspirin by dosing animals with a comparatively higher dose of aspirin (100 mg/kg/d) relative to naproxen sodium (10.9 mg/kg/d) – at least 3 fold higher when considering maximum recommended human doses. Nonetheless, our results are consistent with previous studies which found that only very high doses of aspirin were able to inhibit bone healing in rats and rabbits [41,42]. Furthermore, one group has reported that administration of aspirin was beneficial for fracture healing in osteoporotic rats [43]. As a result, more study into the specific mechanism of action in bone downstream of aspirin and its metabolites may reveal potential therapeutic targets for relieving bone pain without negatively affecting the skeleton.

It is important to note that the overall effects of chronic NSAID usage on general skeletal health, independent of loading, remain poorly understood. Early reports suggested a mild beneficial effect of regular NSAID usage on the skeleton, including reduced measures of bone resorption and increased BMD [44,45]. However, other studies found no effect, dose-dependent effects, or negative effects on fracture risk and BMD due to regular NSAID usage [46,47]. Although the preponderance of clinical and preclinical data suggests that NSAID usage is deleterious for fracture repair [48], clinical studies are contradictory with regards to general bone healing (including spinal fusion); the literature includes roughly equal numbers of recommendations that NSAIDs should be either avoided or prescribed [49]. Furthermore, there is a distinct lack of clinical evidence regarding the effects of NSAIDs on stress fracture healing and only a few preclinical studies [50]. In one study, both selective and non-selective NSAIDs were observed to delay ulnar stress fracture healing in rats [51]. In that study, the authors did not observe any changes in woven bone area with or without normalization to cortical bone area at 2, 4, or 6 weeks after stress fracture as quantified using dynamic histomorphometry. In contrast, we observed a significant decrease in woven bone volume 7 days after stress fracture in naproxen treated mice by microCT quantification. Our result is generally consistent with the only other preclinical study reporting the effects of NSAIDs on stress fracture repair, in which the authors observed that celecoxib significantly decreased bone formation rate during repair [52]. Although it is difficult to directly compare the results given the number of distinct laboratory conditions (animal model, time point, measurement technique), these studies are generally consistent with our conclusion that NSAIDs negatively affect stress fracture repair.

Despite potential contraindications, NSAIDs are often recommended to patients for pain relief to avoid dangerous and potentially addictive prescription narcotics [53]. However, widespread therapeutic use of anti-NGF antibodies, a new class of non-opioid pain relievers, may be imminent [54-56]. In initial preclinical work, anti-NGF antibodies were found to be highly effective for relieving bone pain, including osteoarthritis, fracture, and cancer-related pain [31,57]. Moreover, the humanized monoclonal anti-NGF antibody Tanezumab (Pfizer & Lilly) received FDA Fast Track approval in 2017 for the treatment of chronic osteoarthritis and low back pain. Importantly, the phase III clinical trials for this medication were temporarily halted in 2010 due to an increased incidence of adverse events, particularly in patients also taking NSAIDs [58]. In our study, we found that both aspirin and naproxen significantly decreased Ngf expression 3 hours after a stress fracture. More study is required to determine if this effect is downstream of COX2 inhibition or an alternative effect of these particular NSAIDs (or their metabolites). Nonetheless, these data suggest that concurrent exposure to NSAIDs and anti-NGF antibodies will result in a strong decrease in NGF-TrkA signaling in bone, which may be potentially deleterious.

## EXPERIMENTAL PROCEDURES

### Mice

All procedures involving mice were approved by the Institutional Animal Care of Use Committee of Thomas Jefferson University (protocol #01919). Female C57BL/6J mice were obtained from the Jackson Laboratory (Stock #000664) at 15 weeks of age, and allowed to acclimate in our animal facility for one week prior to experimentation.

### NSAIDs

Mice were randomly separated into three groups: aspirin, naproxen, and control. Aspirin and naproxen treatments were prepared by dissolving acetylsalicylic acid (Sigma, A5376) and naproxen sodium (Acros Organics, AC43671) in ddH2O to final concentrations of 624 mg/L and 68.0 mg/L, respectively. Based on an expected intake of approximately 4 mL per day per mouse, the average daily dose is 100 mg/kg and 10.9 mg/kg, respectively. These dosages are approximately 13% (aspirin) and 4% (naproxen sodium) of the maximum recommended human dose for a 65 kg individual, as determined by the conversion guidelines provided by the FDA Center for Drug Evaluation and Research. Medications were provided 16-24 hours before the first bout of loading and replenished every 2-3 days.

### Mechanical Loading

Each mouse was subjected to mechanical loading intended to induce either lamellar bone formation (LBF) or woven bone formation (WBF) using standard axial forelimb compression [59]. Before loading, all mice received buprenorphine (0.12 mg/kg, IP) and were anesthetized using isoflurane (2-3%, inhaled) for the duration of loading. Deep anesthesia was confirmed by hindpaw pinch before securing the right forelimb in specially designed fixtures for loading using a material testing system (TA Instruments Electroforce 3200 Series III). For LBF loading, a 0.3 N compressive preload was followed by a 2 Hz rest-inserted sinusoidal waveform with a peak force of 3.0 N for 100 cycles. Mice received six bouts of LBF loading over a period of two weeks. The loading parameters for WBF loading were selected as a function of ultimate force and total displacement to fatigue fracture. First, ultimate force was determined using monotonic compression to failure by displacement ramp (0.1 mm/s). Next, total displacement to fatigue fracture was determined using a sinusoidal waveform of 3.6 N (75% of the ultimate force) at 2 Hz. Thus, for WBF loading of experimental animals, a 0.3 N compressive preload was applied followed by a cyclic sinusoidal waveform of 3.6 N at 2 Hz until 66% of the total displacement for fatigue fracture (0.84 mm), relative to the 10th cycle, was achieved. In both cases, mice were placed back in their cages and allowed unrestricted activity following loading.

### Dynamic Histomorphometry

To quantify bone formation rates following LBF loading, calcein (10 mg/kg) and alizarin red (30 mg/kg) were administered on days 3 and 10 after the first bout of loading. Both forelimbs were harvested and fixed overnight in neutral buffered formalin (NBF), dehydrated in increasing grades of ethanol, and embedded in polymethylmethacrylate. Sections were cut at 100μm from the mid-diaphysis of the ulna using a low-speed saw, then mounted with Eukitt, and polished to a thickness of about 50 μm. Images were captured using fluorescence microscopy (Nikon Eclipse E600) and analyzed for endosteal (Es) and periosteal (Ps) mineralizing surface (MS/BS), mineral apposition rate (MAR), and bone formation rate (BFR/BS) as defined by the ASBMR Committee for Histomorphometry Nomenclature [60].

### MicroCT

Bones to be analyzed by microCT were harvested, fixed overnight in NBF, then stored in 70% EtOH until analysis. Each bone was scanned using a Bruker Skyscan 1275 microCT system equipped with a 1 mm aluminum filter. Forelimbs and femurs were scanned using 55 kV and 181 μA with a 78 ms exposure time, whereas tibiae were scanned at 65 kV and 153 μA with a 62 ms exposure time. Transverse scan slices were obtained by placing the long axis of the bone parallel to the z axis of the scanner, using an isometric voxel size of either 8 or 12 micron. Images were reconstructed using nRecon (Bruker) and analyzed using CTan (Bruker).

### Three-point Bending

Bones to be analyzed for three-point bending were harvested and immediately stored at −20 °C in PBS-soaked gauze. Femurs were thawed, then scanned with microCT before performing mechanical testing. For standard three-point bending, each femur was oriented on a standard fixture with femoral condyles facing down and a bending span of 8.2 mm. For notched three-point bending, the notch was created on the posterior surface of the femoral mid-diaphysis with a low speed diamond wafering blade under hydrated conditions. The notch was sharpened by hand with a razor blade applied with 1 μm diamond polish. In either case, a monotonic displacement ramp of 0.1 mm/s was applied until failure, with force and displacement acquired digitally. The force-displacement curves were converted to stress-strain using microCT-based geometry and analyzed using a custom GNU Octave script.

### ELISA assays

Immediately following euthanasia, blood was harvested via cardiac puncture and allowed to coagulate for 30 minutes before centrifugation at 5000 g for 10 minutes to isolate serum, which was stored at −80°C before use. Serum levels of receptor activator of nuclear factor kappa-B ligand (RANKL) and osteoprotegerin (OPG) were quantified via enzyme-linked immunosorbent assays (Quantikine^®^ ELISA, R&D Systems). Serum was diluted 2-fold for RANKL and 5-fold for OPG. Triplicates of 50μL per diluted sample were assayed. Results were compared to a linear standard curve. Similarly, mouse prostaglandin E2 (PGE2) levels in the serum were measured via ELISA (CUSABIO). Samples were diluted 2-fold, and duplicates of 50μL per sample were assayed. A four parameter logistic standard curve was created using Curve Expert (Hyams Development).

### Hydroxyproline Assay

Collagen content was indirectly quantified in the bone and serum using a hydroxyproline assay kit (Sigma), according to the manufacturer’s instructions. Briefly, the tibial sample was decalcified in 14% EDTA for 7 days, then placed in a 1:2 chloroform:methanol solution for 24 hours to extract lipids.

Finally, bone samples were lyophilized overnight. Next, either bone samples or 100 μL of serum was hydrolyzed in 100 μL of 12 N HCl overnight at 120°C. After adding approximately 5 mg of activated charcoal, the samples were centrifuged at 10000 g for 5 minutes and the supernatant was transferred to a new tube. For bone, samples were diluted 100-fold in ddH2O. Finally, 10 uL of each sample was analyzed in triplicate.

### Collagen Crosslinking

Collagen crosslinking was assessed by pepsin digestion and sodium dodecyl sulfate polyacrylamide gel electrophoresis (SDS-PAGE). 5 mm of the tibial mid-diaphysis was decalcified in 14% EDTA, then homogenized in 200 μL of 0.5 M acetic acid on ice. Next, 50 μL of 25 mg/mL pepsin was added before samples were placed on a rotator at 4 °C for 24 hours. Finally, 30μL of each sample was run on 6% separating gel.

### Picrosirius Red Staining and Second Harmonic Generation Imaging

Fibrillar collagen was quantitatively assessed using picrosirius red staining followed by second harmonic generation (SHG) imaging. First, the femoral mid-diaphysis was isolated and decalcified in 14% EDTA for 7 days. After embedding in paraffin, 4 μm thick sections were generated and stained for 1 hour, according to the manufacturer’s instructions (Polysciences, #24901). Finally, sections were mounted on microscope slides with Eukitt and imaged with a circularly polarized light microscope (Nikon Eclipse LV100POL). Best-fit color thresholds were determined by individual pixel selection, and each pixel was categorized to red, yellow, or green (NIS-Elements AR 4.5.00). Each section was subsequently imaged using a two-photon microscope equipped with a 25X water immersion objective (Olympus FV1000MPE). The sections were imaged at 860 nm with 5% laser power, then quantified using FIJI [61].

### Behavioral Testing

To determine analgesic efficacy, forelimb asymmetry tests were performed on WBF-loaded mice on days 0 and 3 to 7. Mice were placed inside a clear, cylindrical tube and video recorded for five minutes. Two mirrors were positioned at an angle that reflected the backside of the tube. For each independent incidence of vertical exploration in which mice stood on their hindlimbs and supported their body weight by placing one or both forepaws on the tube, a score of 1 was given if only the right forepaw was used; 0.5 was given if both forepaws were used; 0 was given if only the left forepaw was used, as in previous studies [62]. A minimum of 10 and up to 42 independent incidents of exploration per mouse were compared to pre-injury baseline.

### Quantitative RT-PCR

After harvesting forelimbs, the proximal and distal ends of the bone were cut off and the marrow was removed by brief centrifugation at 13000 g before placing into TRIzol (Ambion). After pulverization in liquid nitrogen (SpexMill 6750), total RNA was collected according to the manufacturer’s protocol. Next, RNA (0.5 μg) was reversely transcribed using iScript Reverse Transcription Supermix (Bio-Rad). Finally, cDNA (1.6 μL) was amplified using PowerUp SYBR Green Master Mix (Applied Biosystems) under standard PCR conditions. Samples were run in triplicate and normalized to GAPDH expression using the ddCT method. Primer sequences (Table S2) were designed by using Primer-BLAST (NCBI).

### Statistics

All results are presented as mean ± standard error, unless otherwise noted. Statistical analyses were performed in Prism 7 (GraphPad) using ordinary one-way or two-way ANOVA with Dunnett’s correction for multiple comparisons. An adjusted p-value of less than 0.05 was considered significant.

## ACKNOWLEDGEMENTS

The authors thank Drs. Julie Hughes and R. Wayne Matheny for helpful conversations and Drs. Jola Fertala and Andrzej Steplewski for technical assistance. This study was supported by startup funds provided by the Rothman Institute.

## AUTHOR CONTRIBUTIONS

Conceptualization, R.E.T.; Investigation, J.P., A.F., and R.E.T.; Writing, J.P., A.F., and R.E.T.; Supervision, R.E.T. The authors have no conflict of interest.

## Supplemental Information

**Table S1.** Naproxen treatment prevents load-induced bone formation. Values are presented as mean ± standard deviation. * p < 0.05 vs. Non-Loaded. n = 6-7 per group. **Related to Figure 1.**

| Vehicle |  |  |  |  |  |  |
| --- | --- | --- | --- | --- | --- | --- |
| | PS.MS/BS (%) | Es.MS/BS (%) | Ps.MAR ($\mu\text{m}/\text{day}$) | Es.MAR ($\mu\text{m}/\text{day}$) | Ps.BFR/BS ($\mu\text{m}^3/\mu\text{m}^2/\text{day}$) | Es.BFR/BS ($\mu\text{m}^3/\mu\text{m}^2/\text{day}$) |
| **Loaded** | $0.32 \pm 0.07$ * | $0.61 \pm 0.20$ * | $0.86 \pm 0.24$ * | $1.60 \pm 0.57$ * | $0.28 \pm 0.11$ * | $1.12 \pm 0.72$ * |
| **Non-Loaded** | $0.08 \pm 0.04$ | $0.36 \pm 0.22$ | $0.37 \pm 0.16$ | $0.75 \pm 0.58$ | $0.03 \pm 0.02$ | $0.38 \pm 0.58$ |
| Aspirin |  |  |  |  |  |  |
| | PS.MS/BS (%) | Es.MS/BS (%) | Ps.MAR ($\mu\text{m}/\text{day}$) | Es.MAR ($\mu\text{m}/\text{day}$) | Ps.BFR/BS ($\mu\text{m}^3/\mu\text{m}^2/\text{day}$) | Es.BFR/BS ($\mu\text{m}^3/\mu\text{m}^2/\text{day}$) |
| **Loaded** | $0.28 \pm 0.09$ * | $0.40 \pm 0.13$ | $0.67 \pm 0.17$ * | $1.11 \pm 0.60$ | $0.20 \pm 0.09$ * | $0.50 \pm 0.34$ |
| **Non-Loaded** | $0.07 \pm 0.03$ | $0.25 \pm 0.04$ | $0.31 \pm 0.20$ | $0.35 \pm 0.26$ | $0.03 \pm 0.02$ | $0.09 \pm 0.07$ |
| Naproxen |  |  |  |  |  |  |
| | PS.MS/BS (%) | Es.MS/BS (%) | Ps.MAR ($\mu\text{m}/\text{day}$) | Es.MAR ($\mu\text{m}/\text{day}$) | Ps.BFR/BS ($\mu\text{m}^3/\mu\text{m}^2/\text{day}$) | Es.BFR/BS ($\mu\text{m}^3/\mu\text{m}^2/\text{day}$) |
| **Loaded** | $0.18 \pm 0.07$ | $0.37 \pm 0.15$ | $0.54 \pm 0.23$ | $0.69 \pm 0.30$ | $0.10 \pm 0.07$ | $0.27 \pm 0.19$ |
| **Non-Loaded** | $0.10 \pm 0.07$ | $0.23 \pm 0.07$ | $0.43 \pm 0.14$ | $0.18 \pm 0.12$ | $0.04 \pm 0.04$ | $0.05 \pm 0.04$ |

**Figure S1.**
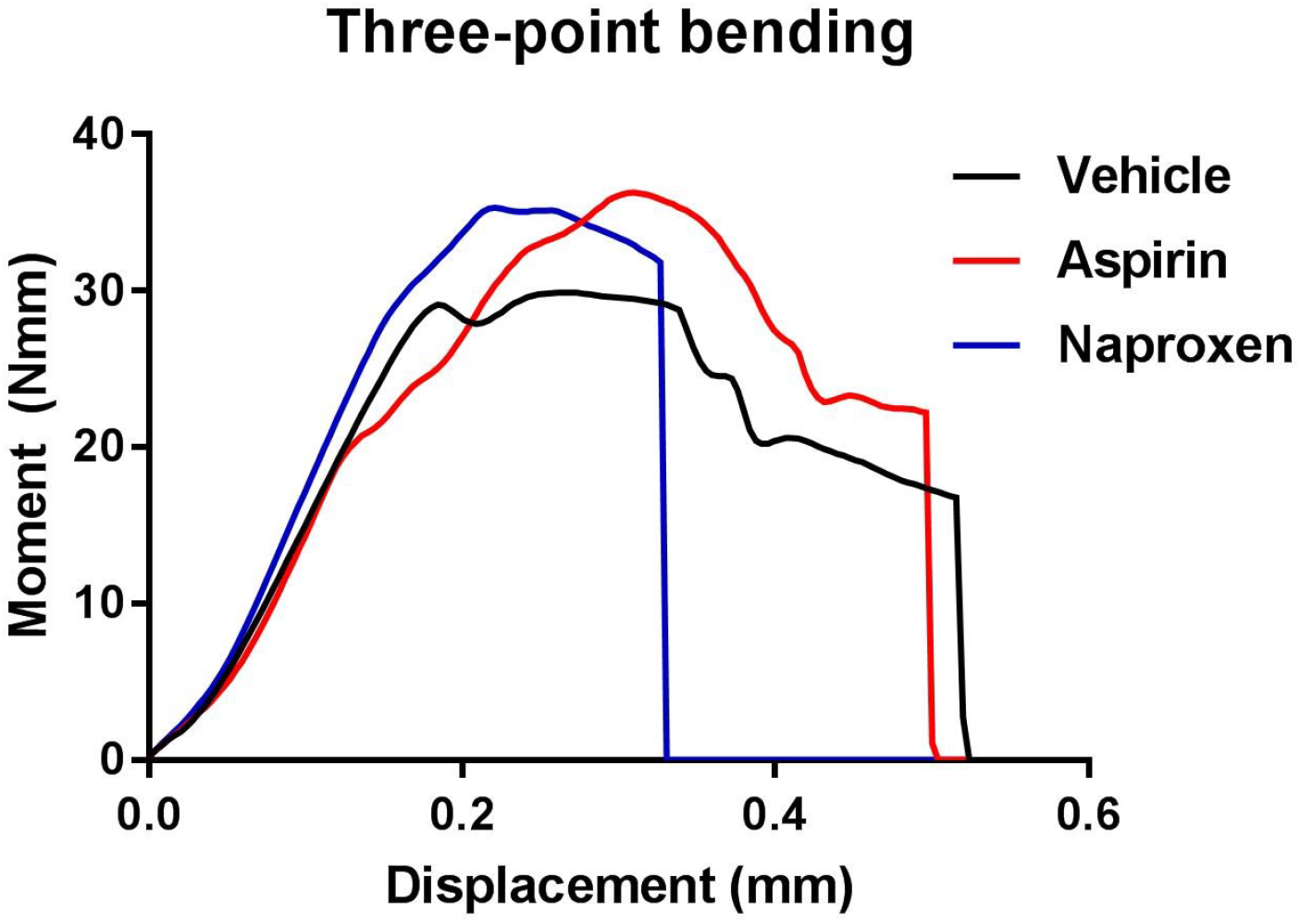
Naproxen treatment alters post-yield behavior. These representative moment-displacement curves from femoral three-point bending illustrate the similar pre-yield behavior and altered post-yield behavior in each treatment group. **Related to Figure 2**.

**Table S2.**
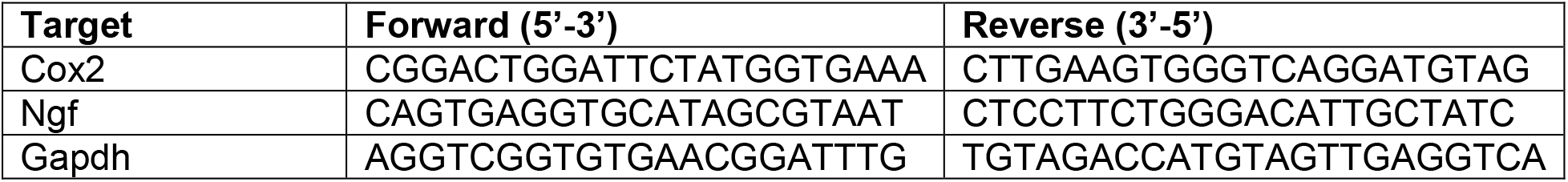
Primers used for qRT-PCR. **Related to Figure 6 Target**

